# Prophage exotoxins enhance colonization fitness in epidemic scarlet fever-causing *Streptococcus pyogenes*

**DOI:** 10.1101/2020.05.17.095000

**Authors:** Stephan Brouwer, Timothy C. Barnett, Diane Ly, Katherine J. Kasper, David M.P. De Oliveira, Tania Rivera-Hernandez, Amanda J. Cork, Liam McIntyre, Magnus G. Jespersen, Johanna Richter, Benjamin L. Schulz, Gordon Dougan, Victor Nizet, Kwok-Yung Yuen, Yuanhai You, John K. McCormick, Martina L. Sanderson-Smith, Mark R. Davies, Mark J. Walker

## Abstract

The re-emergence of scarlet fever poses a new global public health threat. The capacity of North-East Asian serotype M12 (*emm*12) *Streptococcus pyogenes* (group A *Streptococcus*, GAS) to cause scarlet fever has been linked epidemiologically to the presence of novel prophages, including prophage ΦHKU.vir encoding the secreted superantigens SSA and SpeC and the DNase Spd1. Here we report the comprehensive molecular characterization of ΦHKU.vir-encoded exotoxins. We demonstrate that streptolysin O (SLO)-induced glutathione efflux from host cellular stores is a previously unappreciated GAS virulence mechanism that promotes SSA release and activity, representing the first description of a thiol-activated bacterial superantigen. Spd1 is required for optimal growth in human blood, confers resistance to neutrophil killing, and degrades neutrophil extracellular traps (NETs). Investigating single, double and triple isogenic knockout mutants of the ΦHKU.vir-encoded exotoxins, we find that SpeC and Spd1 act synergistically to facilitate nasopharyngeal colonization in a mouse model. These results offer insight into the etiology and pathogenesis of scarlet fever-causing GAS mediated by phage ΦHKU.vir exotoxins.

## Introduction

Scarlet fever is a superantigen-mediated acute infectious disease caused by the human-adapted pathogen group A *Streptococcus* (GAS). Scarlet fever was a leading cause of death in children in the early 1900s, but its incidence steadily declined during the 20th century^1,2^. Large regional outbreaks of scarlet fever re-emerged in North-East Asia in 2011, and the United Kingdom in 2014^3-10^, with factors driving disease resurgence remaining a mystery. Alarmingly, recent studies report GAS outbreak strains in other countries^11-13^, heightening the need for global surveillance^14^.

Potential triggers for these new scarlet fever epidemics remain unclear, but accumulating epidemiological evidence indicates that novel prophages and antibiotic resistance elements have played a significant role in the evolution, virulence and diversification of scarlet fever causing GAS strains in North-East Asia^4,15-17^. Detailed phylogenetic analyses of GAS outbreak isolates from mainland China and Hong Kong prove that the increase in scarlet fever cases was neither *emm*-type specific nor caused by the spread of a single scarlet fever producing clone. Instead, multiclonal scarlet fever outbreak strains are commonly associated with the acquisition of related exotoxin-carrying mobile genetic elements^15,17^. Prophages encoding combinations of the streptococcal superantigens SSA and SpeC, and the DNase Spd1, appear to play an important role in the evolutionary pathway that lead to the emergence of more virulent strains, particularly in North-East Asia^4-6,15-18^. However, robust evidence defining the mechanistic contribution of prophage-encoded exotoxins to the pathogenesis of scarlet fever is lacking. A universal feature of superantigens is their ability to cross-link major histocompatibility complex II molecules on antigen-presenting cells and the variable region of the β-chain of T cell receptor (TCR). This cross-linkage results in TCR Vβ-specific activation of large populations of human T cells, without prior antigen processing, rendering superantigens the most potent T cell mitogens known to date^19^. Recent studies suggest that such T cell activation contributes to the establishment of GAS infection at mucosal surfaces^20,21^. Here, we investigate the regulation of ΦHKU.vir encoded exotoxin genes *ssa, speC* and *spd1*, and their impact on the virulence of scarlet fever-causing GAS. Exotoxin-driven enhanced colonization provides an evidence-based hypothesis for the reemergence of scarlet fever globally.

## Results

### Regulation of ΦHKU.vir exotoxins

The majority of GAS *emm12* clones from scarlet fever outbreaks in North-East Asia carry superantigens SSA and SpeC and the DNase Spd1, as well as integrative and conjugative elements (ICE) encoding tetracycline (*tetM*) and macrolide (*ermB*) resistance^4,15,17^. Penicillin remains the treatment of choice for GAS pharyngitis. However, in many countries macrolides are commonly used as first-line therapy for upper respiratory tract infections in primary health-care settings^22^. To investigate whether antibiotic treatment stress affects either prophage induction or superantigen expression, macrolide-resistant GAS *emm12* scarlet fever isolate HKU16 harboring ΦHKU.vir and ICE–*emm*12 was grown in THY medium containing erythromycin (2 µg/ml), the recommended drug in patients with penicillin hypersensitivity^23^. RNA-seq analysis showed that erythromycin treatment did not affect the gene expression pattern of ΦHKU.vir (Fig. 1a), whereas expression of ICE-*emm12*-encoded *ermB* gene was significantly increased (supplementary Fig. S1). Mitomycin C, a DNA-damaging agent known to induce GAS prophage^24^, effectively induced ΦHKU.vir housekeeping and structural gene expression (Fig. 1a, supplementary Fig. S1). Similar to prophage-encoded virulence factor cargo genes in *emm3* GAS^24^, mitomycin C did not induce expression of the virulence cargo genes *ssa, speC* and *spd1*, pointing to differential control of exotoxin expression in ΦHKU.vir.

**Figure 1:**
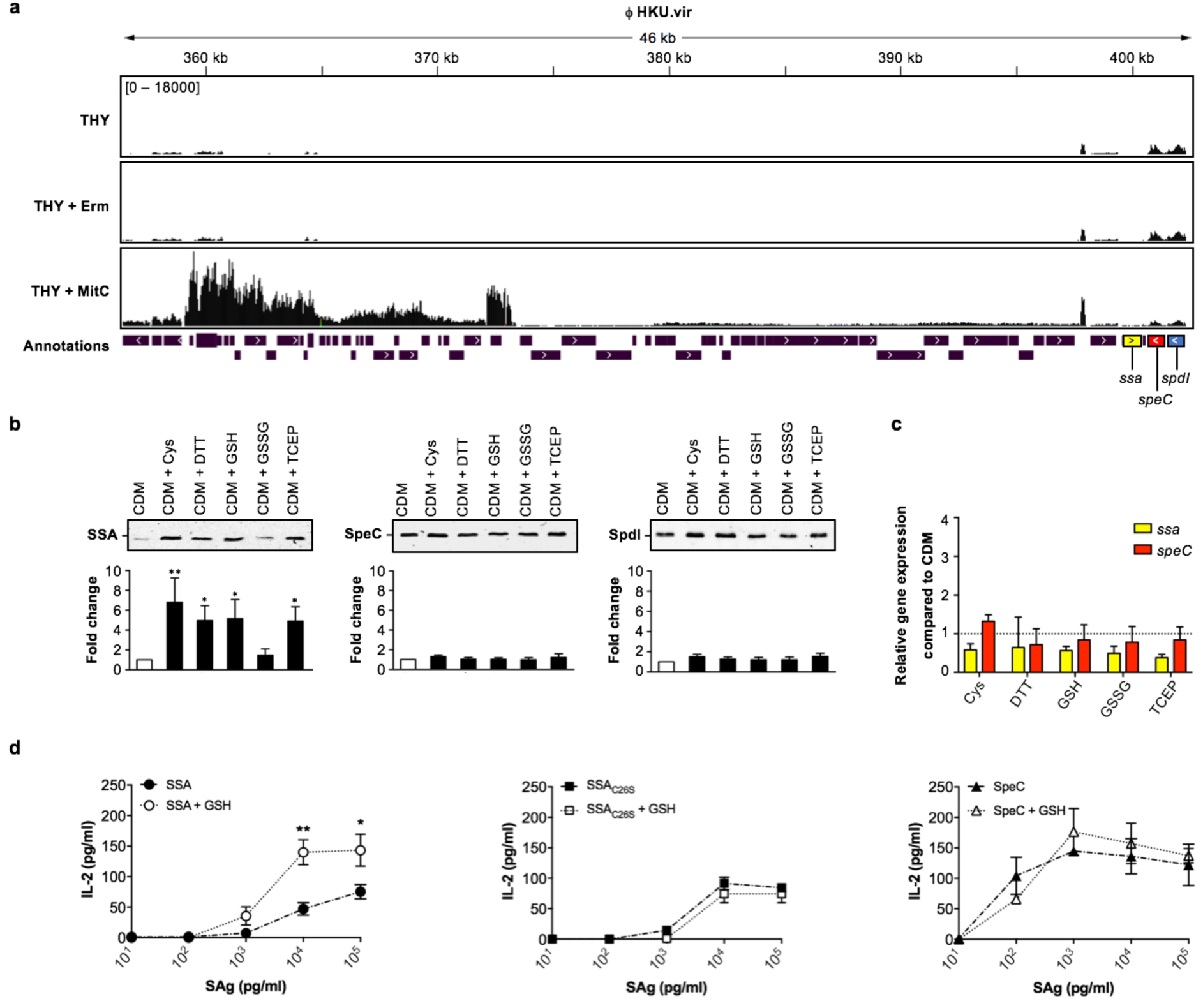
Post-transcriptional thiol-based regulation of SSA. (**a**) RNAseq expression profile of ΦHKU.vir in the macrolide- and tetracycline-resistant GAS *emm12* isolate HKU16, grown in THY broth with sub-inhibitory concentrations of erythromycin (Erm) and mitomycin C (MitC). The plots illustrate the overall coverage distribution displaying the total number of sequenced reads. The region that encodes exotoxin genes (*ssa* in yellow, *speC* in red and *spd1* in blue) is indicated. (**b**) Immunoblot detection of SSA, SpeC and Spd1 in culture supernatants of HKU16 grown in a chemically defined medium (CDM) in the presence of various redox-active compounds. Western blot signal intensities were quantified with ImageJ. Statistical significance is displayed as *p<0.01 and **p<0.001 by one-way ANOVA (n = 4). (**c**) Quantitative real-time PCR of *ssa* and *speC* transcripts in HKU16 grown in CDM treated with 2 mM of the indicated redox-active compounds. (**d**) Superantigen (SAg) activation of human PBMCs (2 × 10^5^ cells per well) with SSA (circular), SSA_C26S_ (square), and SpeC (triangular) at the indicated concentrations in absence (black; dash-dot line) or presence of 2 mM of GSH (white; dotted line), using human IL-2 as a readout. Bars represent the mean ± SEM. Statistical significance is displayed as *p<0.01 and **p<0.001 by Student’s t test.

### Thiol-mediated induction of SSA release

While there is good evidence that phage-associated exotoxins SpeC and Spd1 are induced during host-pathogen interactions^25,26^, comparatively less is known about the control of SSA expression. The *ssa* gene is frequently associated with scarlet fever isolates from North-East Asia^15,17^. As SSA production is detectable upon growth in a chemically-defined medium^16^, we undertook a limited small molecule screen that identified cysteine, but not any other amino acid, as a factor specifically increasing abundance of the exotoxin SSA in culture supernatants (Fig. 1b, supplementary Fig. S2). Cysteine is uniquely chemically reactive, due to its thiol-containing side chain. We therefore examined whether SSA production was subject to thiol-mediated regulation. Both dithiothreitol (DTT) and the reduced form of glutathione (GSH) increased SSA production in GAS culture supernatants (Fig. 1b). In contrast, oxidized glutathione (GSSG) did not enhance SSA levels. Higher SSA production was also detected in GAS cultures treated with thiol-free reducing agent tris(2-carboxyethyl)phosphine (TCEP), suggesting that exposure to reducing conditions enhances SSA production. The levels of secreted SpeC and Spd1 were unaffected by any of these treatments (Fig. 1b). Quantitative real-time PCR of the *ssa* and *speC* transcripts suggested that reducing agents are acting as post-transcriptional enhancers of SSA release (Fig. 1c). To validate the requirement for thiols (reducing conditions) in SSA regulation, we also performed alkylation of cysteine with acrylamide prior to treatment, resulting in significant reduction of SSA, but not SpeC, release (supplementary Fig. S3a).

### SSA is a thiol-activated superantigen

SSA contains a surface-exposed Cys-26 residue that, based on the crystal structure of the homologous SpeA superantigen in complex with TCR Vβ^27^, is predicted to lie within the TCR binding interface (supplementary Fig. S3b). Prior site-directed mutagenesis has revealed a role for Cys-26 in the mitogenic activity of SSA on human T cells by preventing disulphide-linked dimer formation between the surface-exposed Cys-26 residues of SSA^28^. Although a SSA dimer was not detectable in HKU16 culture supernatants (supplementary Fig. S3c), we detected dimer formation by purified recombinant SSA (supplementary Fig. S3d) which led us to investigate possible redox sensitivity of SSA activity. GSH, the major low-molecular-weight thiol in living cells, markedly increased the mitogenic potency of recombinant SSA on human T cells by ∼10-fold as assessed by enhanced IL-2 production (Fig. 1d). However, thiol activation by GSH was absent in SSA carrying a cysteine-to-serine substitution at position 26 (SSA_C26S_), underscoring a critical role for the Cys-26 residue in thiol-mediated activation. In contrast to SSA, the activity of SpeC, one of the most potent T cell mitogens^29^, was unaffected by GSH treatment (Fig. 1d). These data establish a unique role for thiols in SSA regulation and support a model where reducing agents not only increase levels of extracellular SSA monomer, but also directly enhance SSA-mediated T cell stimulation. To our knowledge, this is the first report of a thiol-activated superantigen.

### Streptolysin O (SLO) mediates the rapid release of host intracellular glutathione

Like other species of pathogenic Gram-positive bacteria, GAS produces a cholesterol-dependent cytolysin (CDC), streptolysin O (SLO), that perforates host cell membranes^30^. In contrast to plasma and other extracellular fluids that are low in thiol-based antioxidants, the cytosol of mammalian cells is a highly reducing compartment where thiols are present at high concentration. The most abundant non-protein thiol in mammalian cells is GSH, with intracellular concentrations typically in the millimolar range (∼1-11 mM), compared to extracellular concentrations in the low micromolar range^31^. This GSH concentration differential across the plasma membrane led us to speculate that host cell lysis by SLO, itself subject to thiol activation^32^, could provide extracellular GAS with access to the intracellular GSH pool, altering the redox environment and supporting SSA activation.

To test this hypothesis, we first quantified glutathione release after treatment of whole human blood with increasing concentrations of purified SLO. SLO lysed red blood cells (Fig. 2a), and both haemoglobin and total glutathione (GSH + GSSG) accumulated rapidly in plasma in a dose-dependent manner (Fig. 2a). In the context of live GAS, wildtype (WT) scarlet fever-associated strain HKU16 caused significant red blood cell hemolysis after 4 h growth in human blood (Fig. 2b), paralleled by a significant and substantial release of glutathione into plasma (Fig. 2c). In contrast, an isogenic GAS HKU16Δ*slo* mutant did not induce hemolysis and plasma levels of glutathione were unchanged (Fig. 2b, c).

**Figure 2:**
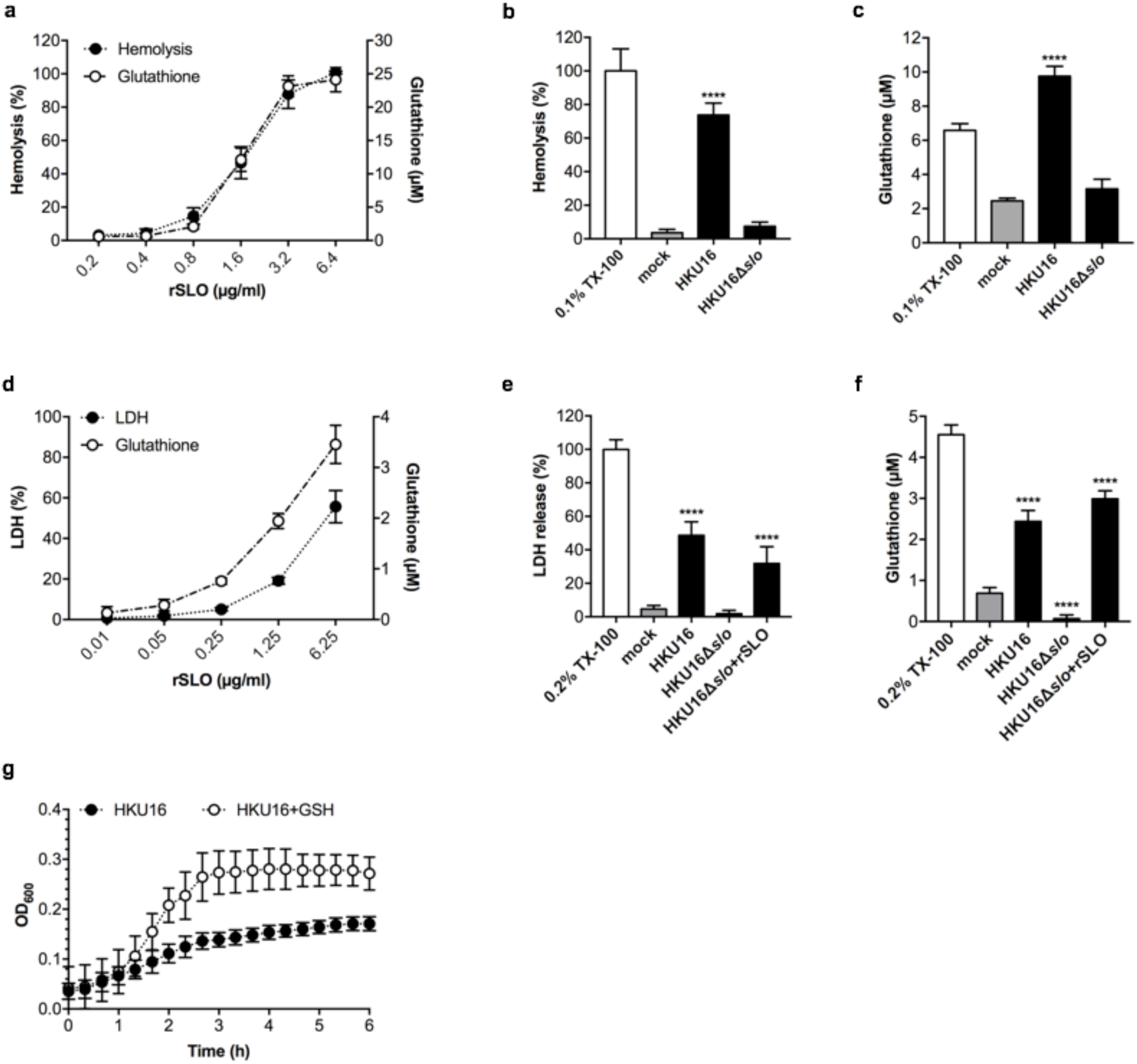
The cytotoxic activity of SLO causes the release of host cytosolic glutathione. (**a**) Dose-dependent hemolytic activity of purified recombinant SLO (rSLO) in whole human blood is accompanied by an extracellular accumulation of glutathione. Hemolysis is expressed as percentage ± SD with respect to the positive control (cells treated with 0.1% Triton X-100 (TX-100)). (**b**) Hemolytic activity of indicated HKU16 strains is expressed as percentage ± SD with respect to the positive control (cells treated with 0.1% TX-100). Blood treated with HBSS (mock) served as a negative control. (**c**) Extracellular accumulation of glutathione in blood infected with indicated HKU16 strains. Blood treated with HBSS (mock) served as a negative control. (**d**) Release of lactate dehydrogenase (LDH) (closed circles) and glutathione (open circles) by D562 cells treated with varying concentrations of recombinant SLO. The release of LDH and glutathione into the culture medium was assessed after 2 h of treatment. LDH release is expressed as percentage ± SD with respect to the positive control (cells treated with 0.2% TX-100). Cells treated with growth medium (mock) served as a negative control. LDH (**e**) and glutathione (**f**) release by D562 cells challenged with indicated HKU16 strains at a multiplicity of infection of 20:1 (bacterial CFU:cells), assessed at 2 h post-infection. Where indicated, rSLO was added to HKU16Δ*slo*-infected cells at a concentration of 6.25 µg/ml. (**g**) Growth curves of HKU16 in D562 cell-free culture medium (EMEM + 10% FBS) supplemented with 2 mM of GSH. Statistical significance is displayed as ****p<0.0001 by one-way ANOVA.

GAS serotype M12 strains belong to *emm* pattern A–C and have been designated as “throat specialists”^33^. In this context, we used human pharyngeal cells (Detroit 562; D562) to study the effect of SLO-induced pore formation on glutathione release as a pharyngitis-relevant cellular model. Lactate-dehydrogenase (LDH) release into the media serves as a marker of host cell membrane integrity. As expected, SLO caused a dose-dependent release of LDH of ∼ 50% at 6.25 µg/ml, confirming disruption of the cell membrane structure (Fig. 2d). Dose-dependent cell death following SLO exposure was again associated with a progressive increase in glutathione level in the media (Fig. 2d), indicating that SLO-induced membrane disruption was sufficient to trigger extracellular release of host cytosolic glutathione stores. Next, levels of LDH and glutathione released by D562 cells following infection with live GAS (multiplicity of infection = 20 bacterial colony forming units:cell) were measured. At 2 h post-infection, WT GAS HKU16 but not the HKU16Δ*slo* mutant induced a significant increase in levels of secreted LDH and glutathione (Fig. 2e, f). The addition of purified pore-forming protein toxin SLO (6.25 µg/ml) to D562 cells grown in the presence of HKU16Δ*slo* markedly elevated extracellular LDH and glutathione to WT HKU16 levels during co-culture. To examine whether the lack of glutathione release following infection with HKU16Δ*slo* (Fig. 2f) might impact other aspects of GAS biology, we measured growth in cell-free medium with and without glutathione supplementation. Supplementation with glutathione strongly promoted growth of WT GAS HKU16 in cell-free medium (Fig. 2g), showing that host-derived glutathione is utilized by GAS for other physiological pathways. Taken together, our data demonstrate that SLO is highly effective at triggering the release of significant amounts of glutathione from host cells, which is utilized for extracellular growth of GAS and likely provides a reducing extracellular microenvironment required for efficient SSA activation *in vivo*.

### ΦHKU.vir-encoded DNase Spd1 enhances fitness of HKU16 in human blood and promotes resistance to neutrophil killing

Horizontal transmission of bacteriophage encoding DNase Sda1/SdaD2 has played a critical role in the emergence and global dissemination of the highly virulent M1T1 clone^34-36^. The phage-encoded DNase Spd1 is linked with the expansion of scarlet fever GAS in North-East Asia^15^. In contrast to Sda1^36^, the contribution of Spd1 to GAS pathogenesis remains largely unexplored, although this nuclease has previously been shown to play a role in nasal shedding in *emm3* GAS^37^.

Unlike the knockout strains HKU16Δ*ssa* and HKU16Δ*speC*, DNase knockout strain HKU16Δ*spd1* showed significantly attenuated growth in human blood (Fig. 3a). Reinforcing these results, complementation of HKU16Δ*spd1* with the WT *spd1* gene (HKU16Δ*spd1*^++^) fully restored growth in human blood (Fig. 3a). Neutrophils are the first immune cell responders to sites of bacterial infection, and thus play a critical role in controlling GAS infection. Examining the role of Spd1 in bacterial susceptibility to human neutrophil killing, knockout strain HKU16Δ*spd1* showed significantly reduced survival compared to the WT and complemented HKU16 strains (Fig. 3b). Formation of web-like lattices composed of chromatin and granular proteins, known as neutrophil extracellular traps (NETs), is a well-established antimicrobial mechanism^38^. Multiple pathogenic microorganisms, including GAS, secrete DNases to dissolve NETs and escape neutrophil mediated killing^39^. To determine the ability of Spd1 to facilitate NET degradation, we used phorbol-myristate acetate (PMA) to induce high levels of NETs from freshly isolated human neutrophils (Fig. 3c) that are sensitive to bovine pancreatic DNase I (Fig. 3c, d). We then incubated PMA-stimulated neutrophils with GFP-expressing GAS for 30 min. NETs exposed to HKU16Δ*spd1* remained intact and covered a significantly greater area in the absence of Spd1 (64.1 ± 3.3 %) compared to NETs infected with WT HKU16 (24.5 ± 4.1 %) and HKU16Δ*spd1*^++^ (21.9 ± 5.2 %) (Fig. 3e, f). Additionally, higher numbers of HKU16Δ*spd1* bacteria were immobilized within NETS compared to WT HKU16 and HKU16Δ*spd1*^++^, while similar levels of NET degradation were displayed by WT HKU16 and HKU16Δ*spd1*^++^ (Fig. 3e, f). These findings demonstrate that Spd1 promotes growth of HKU16 in whole blood, reduces susceptibility to neutrophil mediated killing and facilitates NET degradation.

**Figure 3:**
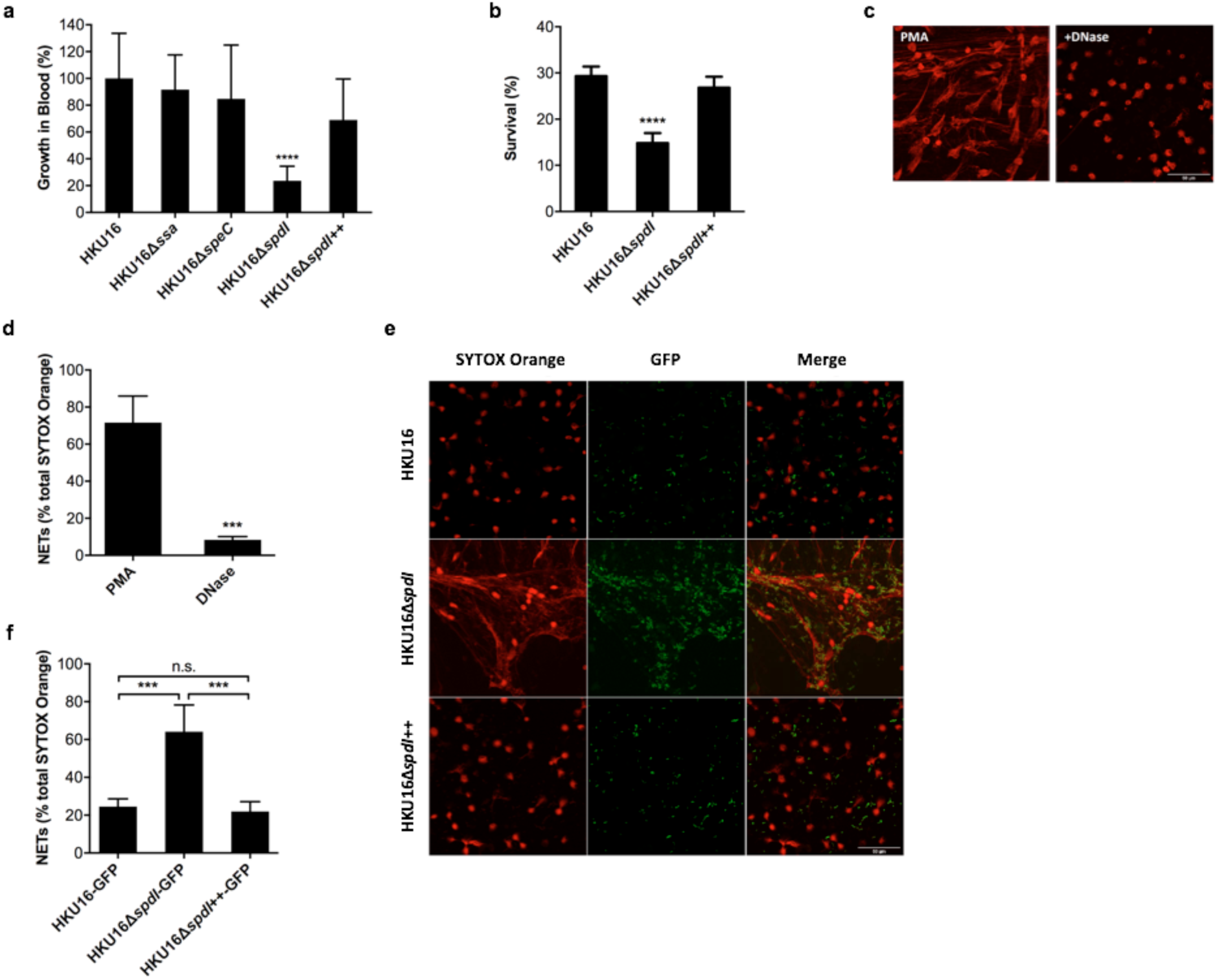
The ΦHKU.vir-encoded DNase Spd1 promotes resistance to neutrophil killing. (**a**) Growth rate of indicated HKU16 strains in whole human blood. Statistical significance is displayed as ****p<0.0001 by one-way ANOVA (**b**) Human neutrophil killing assay. The data represent means ± SEM of six independent experiments. Statistical significance is displayed as ****p<0.0001 by one-way ANOVA (**c**) Purified human neutrophils were stimulated with 25 nM PMA for 3 h to induce neutrophil extracellular traps (NETs). NETs were detected using the extracellular DNA stain SYTOX Orange (red) and images captured using confocal microscopy. Panels show formation of NETs (left) and NET degradation following incubation with bovine pancreatic DNase I as a positive control (right). (**d**) NET quantification of PMA-stimulated neutrophils in the absence or presence of DNase I. Statistical significance is displayed as ***p<0.001 by Student’s t-test. (**e**) Representative images of PMA-stimulated neutrophils following infection with GFP fluorescent GAS (green) for 30 min at a multiplicity of infection of 10 (bacterial CFU:neutrophil). Scale bars represent 50 μm. (**f**) NET quantification of PMA-stimulated neutrophils following incubation with GAS. NET quantification is expressed as a percentage of total SYTOX Orange stained area calculated from a minimum of five randomly selected microscopic fields. Error bars represent means ± SEM from three independent experiments. Statistical significance is displayed as ***p<0.001 by one-way ANOVA.

### Role for ϕHKU.vir-encoded exotoxins in pharyngeal colonization

Previous studies have shown that intranasal infection of mice with GAS can serve as a model to study pharyngeal infection in humans^20,21,40,41^. Humanized mice that express HLA-DR4 and HLA-DQ8 are susceptible to acute nasopharyngeal infection by SpeA-carrying *emm18* GAS^20,21^. To evaluate the role of HKU16 exotoxins in nasopharyngeal infection, we investigated the ability of WT and isogenic mutants to colonize the nasopharynx of HLA-B6 mice transgenic for human CD4 and HLA-DR4-DQ8 genes^42^. The growth phenotype of all single, double and triple HKU16 isogenic knockout mutants (Fig. 4a) was indistinguishable from the parental WT strain (Fig. 4b) and all mutant strains were defective for production of the targeted exotoxins SSA, SpeC, Spd1 and SLO (Fig. c). HLA-B6 mice were infected intranasally with WT HKU16 or isogenic mutants. At 48 h post-infection, significantly fewer bacterial colony forming units (CFUs) were recovered from the complete nasal turbinates of mice infected with HKU16Δ*speC/spd1* compared to WT HKU16 (Fig. 4d). Single isogenic mutant strains of ΦHKU.vir-encoded exotoxins did not show reduced colonization efficiency, suggesting that SpeC and Spd1 act synergistically to enhance nasopharyngeal infection, nor did the additional knockout of the *ssa* gene in the triple mutant strain HKU16Δ*ssa/speC/spd1* further reduce colonization. The attenuated virulence phenotype of HKU16Δ*ssa/speC/spd1* could be fully restored by genetic complementation with WT *ssa, speC* and *spd1* genes (HKU16Δ*ssa/speC/spd1*^++^) (Fig. 4d). Significantly fewer bacterial CFUs were also recovered from HKU16Δ*slo* infected mice (Fig. 4d), confirming the importance of SLO for GAS pathogenicity as demonstrated in previous studies^43-45^.

**Figure 4:**
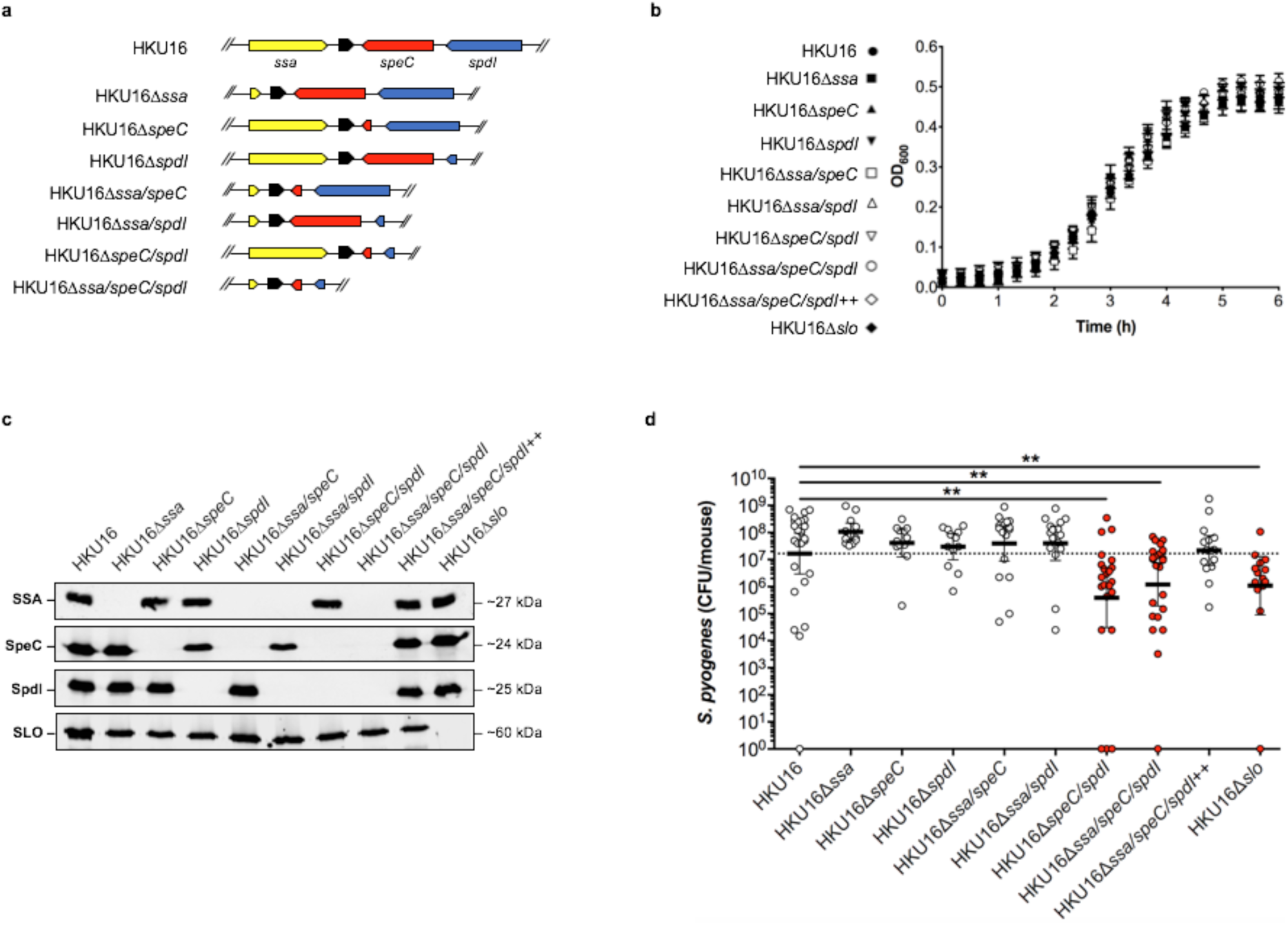
ΦHKU.vir-encoded exotoxins and SLO are critical for HKU16 nasopharyngeal infection. (**a**) Illustration of the genetic in-frame deletions of ΦHKU.vir-encoded exotoxins in HKU16 as described in Materials and Methods. (**b**) Growth curves of indicated HKU16 strains in CDM. (**c**) Immunoblot detection of SSA, SpeC, Spd1 and SLO expression from indicated HKU16 strains. The molecular mass of each protein (kDa) is indicated to the right. (**d**) Individual ‘humanized’ B6 mice that express HLA-DR4, HLA-DQ8 and CD4 were nasally inoculated with ∼1 × 10^8^ bacterial colony forming units (CFU) with indicated HKU16 strains and nasopharyngeal CFUs were assessed at 48 h post-infection. Each symbol represents CFUs from an individual mouse (n ≥ 12). Presented is the geometric mean with 95% confidence interval. Significance was calculated using the Kruskal-Wallis test with the Dunn’s multiple comparisons post-hoc test (**p<0.001).

## Discussion

Mainland China and Hong Kong have witnessed an ongoing outbreak of scarlet fever with ∼500,000 reported cases since 2011^4,14,15,17,46-49^. Alarmingly, case numbers have again significantly increased in recent years posing a heightened global threat to public health (supplementary Fig. S4)^12^. Previous epidemiological surveillance studies have shown that *emm12* is the most prevalent GAS *emm* genotype in clinical cases of scarlet fever in this region^4,15,17^. In contrast with the United Kingdom epidemic, the expansion of scarlet fever– associated *emm12* lineages in North-East Asia has been directly linked to acquisition of two genetic elements: the *tetM*- and *ermB*-carrying multidrug resistance element ICE-*emm12* (and its derivatives) and the prophage ΦHKU.vir, encoding SSA, SpeC and the DNase Spd1^4,15,50^. Consistent with these prior studies, the results presented here demonstrate a direct contribution of ΦHKU.vir acquisition to virulence phenotypes of the scarlet fever-causing *emm12* reference strain HKU16. Using defined genetic knockouts, our data suggest that SpeC and the DNase Spd1 function synergistically to mediate nasopharyngeal colonization, offering an explanation as to why these genes form a conserved genetic module in a variety of distinct GAS prophage^26^.

We also present new insight into the activation of the scarlet fever-associated superantigen SSA, which we reveal as a thiol-activated superantigen. By providing a mechanistic framework of how extracellular GAS gains access to highly abundant intracellular glutathione *in vivo*, we highlight the relationship between SLO-mediated membrane disruption and SSA activity (Fig. 5). Data presented here extend previous studies showing that epithelial cell damage by SLO augments superantigen penetration, which allows for better interaction of superantigens with antigen presenting cells in underlying tissues^51^. Together, these studies suggest that SLO pore formation promotes SSA activation, which may be an important driver in diseases associated with superantigen production, including scarlet fever.

**Figure 5:**
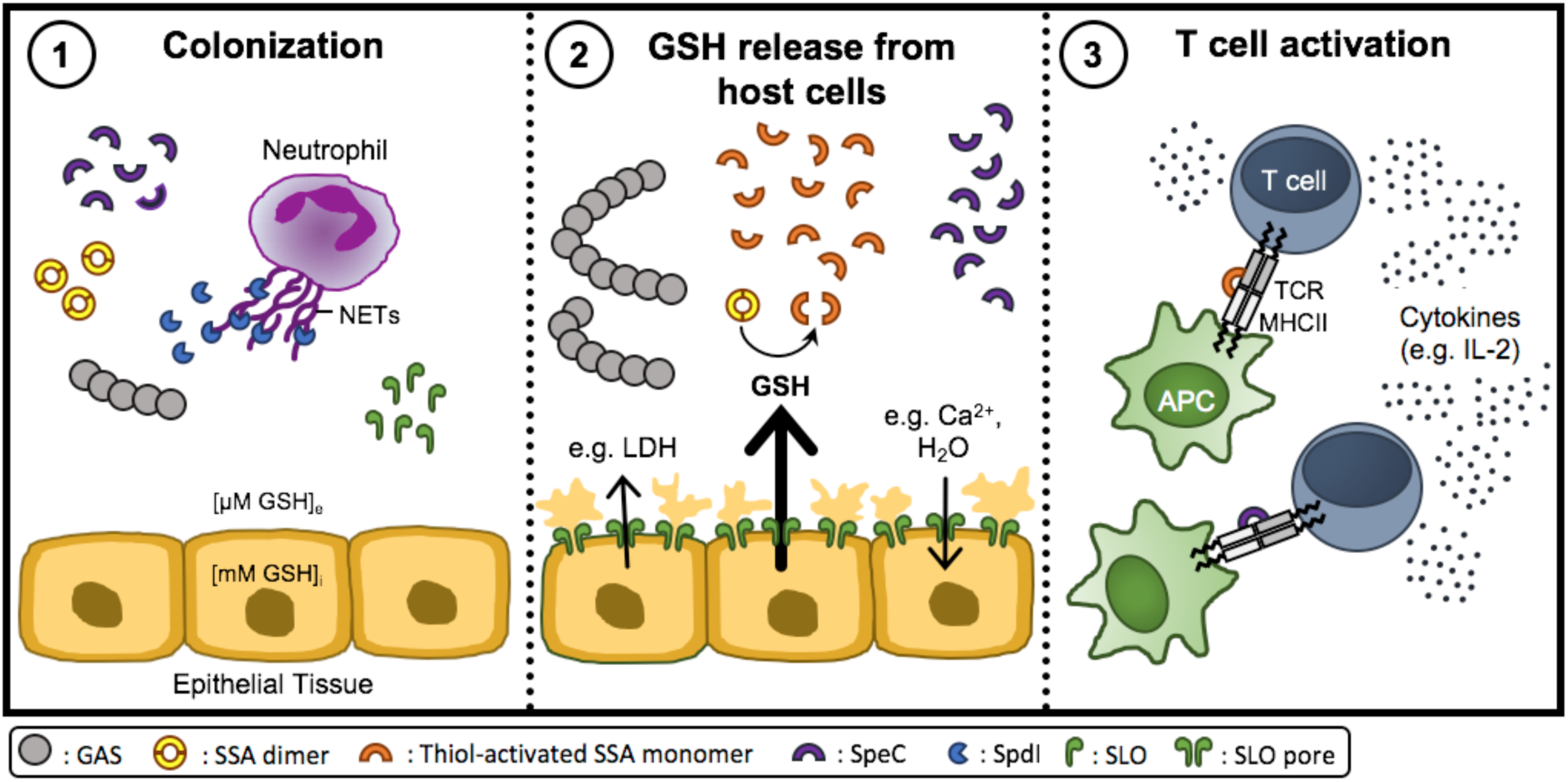
Proposed mechanistic model outlining inter-relationships between SLO mediated cytotoxicity towards epithelial cells and SSA superantigen potency. (**1**) During initial bacterial colonization, GAS secretes the DNase Spd1 to escape neutrophil clearance, allowing GAS to establish infection. (**2**) As infection progresses, SLO binds to host cell membranes and then oligomerizes to form large pores which induces the release of lactate dehydrogenase (LDH) and GSH from perforated host cells as well as cation influx^65,66^. In contrast to cations, glutathione exists at a much higher concentration in the intracellular compartment than the extracellular space (∼1000-fold) causing a significant difference in redox potential across the plasma membrane of eukaryotic cells. This gradient makes the extracellular and intracellular areas, respectively, oxidative and reductive. GSH efflux from perforated cells serves as a stimulus for SSA release, reduces SSA dimers and activates SSA monomers. (**3**) Thiol-activated SSA, in conjunction with other superantigens like SpeC, then crosslinks major histocompatibility complex II molecules on antigen-presenting cells (APCs) and the variable region of the β-chain of T cell receptor (TCR) to induce an overwhelming T cell response with uncontrolled cytokine release.

GAS encodes several glutathione-dependent proteins, yet the bacterium lacks genes for *de novo* glutathione synthesis. This paradox raises the possibility that GAS may coordinate a range of virulence factors through SLO-mediated GSH release. One such factor is glutathione peroxidase (GpoA)^52^, which plays a role in the adaptation of GAS to oxidative stress during inflammation following systemic infection^53^. Microbial acquisition of nutrients *in vivo* is a fundamental aspect of infectious disease pathogenesis, and intracellular bacterial pathogens capitalize on the ubiquitous and highly abundant cytosolic antioxidant GSH^54,55^. Our data support a hypothesis in which extracellular bacterial pathogens such as GAS may have evolved a mechanism to target and hijack host cytosolic glutathione, consistent with the absence of glutathione biosynthetic genes in the GAS genome. While a precise role for SSA in virulence was not conclusively established in the HLA-B6 mouse model, this work exemplifies an interconnected action of GAS virulence determinants such as SLO and SSA, opening new avenues to understand the evolution and emergence of pathogenic clones. As multiple bacterial pathogens encode functional homologs of SLO^30^, glutathione release by cholesterol-dependent cytolysins may constitute a generalized mechanism used by pathogenic bacteria to modulate their physiological response to host cells, including through the post-transcriptional activation of virulence-associated proteins.

Our findings show that GAS HKU16 requires the ΦHKU.vir-encoded exotoxins SpeC and Spd1, and SLO, to efficiently colonize the HLA-B6 mouse model. We hypothesise that prophage-encoded exotoxin acquisition has enhanced colonization fitness of scarlet fever-causing GAS *emm12* clones in North-East Asia. The atypical presence of genes encoding superantigens such as SSA in *emm12* isolates^56^ could provide a framework allowing for clonal expansion of GAS in a naïve population. The spread of such prophage-containing GAS is therefore of great public health concern and heightened efforts are needed to instigate global surveillance systems. Recent evidence of interspecies transfer of *speC*- and *spd1*-containing prophage in the US should serve as a warning for the dissemination of these virulence-enhancing genes into other pathogenic streptococci^57^.

## Acknowledgments

We kindly acknowledge James McCluskey (University of Melbourne, Melbourne, Australia) for providing HLA-B6 mice transgenic for human CD4 and HLA-DR4-DQ8 genes; Luke McAlary (University of Wollongong, Wollongong, Australia) for assistance with microscopy; and assistance from the sequencing and pathogen informatics core teams at The Wellcome Trust Sanger Institute (Hinxton, UK). We thank Thomas Proft, Mitchell Acev, Heema Vyas and Jason McArthur for providing genetic constructs. This work was supported by grants from the National Health and Medical Research Council of Australia, The Wellcome Trust, UK, the National Institutes of Health, USA, and the Canadian Institutes of Health Research.

## Author contributions

SB, TCB, GD, VN, KYY, YY, JKM, MLSS, MRD and MJW conceived this study. SB, TCB, DL, KJK, BLS, GD, VN, KYY, YY, JKM, MLSS, MRD and MJW planned experiments. SB, TCB, DL, KJK, LM, MGJ, DMPDO, TRH, AJC, JR and MRD performed experiments. GD, VN, KYY and JKM provided essential reagents and strains. SB and MJW wrote the manuscript and all authors reviewed and revised the manuscript.

## Methods

### Bacterial strains, growth conditions and construction of HKU16 mutant strains

The *emm12* GAS scarlet fever isolate HKU16^4^ and isogenic derivatives were routinely grown at 37°C on 5% horse blood agar or statically in Todd-Hewitt broth supplemented with 1% yeast extract (THY) or chemically defined medium (CDM; Gibco RPMI 1640 with L-glutamine and phenol red (Life Technologies; 11875-093) supplemented with 0.7% (w/v) D-Glucose, 1% (v/v) BME vitamins (Sigma; B6891), 0.15 mM nucleobases (adenine, guanine, and uracil), and 0.02 mM HEPES, pH 7.4). To facilitate fluorescent microscopy experiments, GAS strains were transformed with GFP-expressing plasmid pLZ12Km2-P23R-TA:GFP (supplementary information). *Escherichia coli* (*E. coli*) strains MC1061 or XL1-blue, and BL21(DE3), were used for cloning and protein expression, respectively. *E. coli* was grown in Luria-Bertani medium (LB). Where required, spectinomycin was used at 100 µg/ml (both GAS and *E. coli*), ampicillin was used at 100 µg/ml (*E. coli*), and kanamycin was used at 50 µg/ml (*E. coli*). All bacterial strains and plasmids are listed in Table S1. Isogenic HKU16 mutants were generated using a highly-efficient plasmid (pLZts) for creating markerless isogenic mutants^58^. All PCR primer sequences are provided in Table S1. All gene deletions were confirmed by DNA sequence analysis (Australian Equine Genome Research Centre, University of Queensland, Brisbane, Australia). To examine fitness of WT and mutant strains, GAS were firstly grown overnight on horse blood agar. GAS were then inoculated into CDM to an optical density at 600 nm (OD600) of 0.01. Late-exponential phase GAS grown in CDM (OD600 of 0.4) were resuspended in ATCC Eagle’s Minimum Essential Medium (EMEM; ATCC302003) supplemented with 10% heat-inactivated fetal bovine serum (FBS). Bacteria were then inoculated into 96-well microtiter plates and the growth curves measured using the FLUOstar Omega Microplate Reader (BMG Labtech) at 37°C.

### Transcriptomic analysis and quantitative gene expression studies

Total RNA was routinely isolated from bacterial cells as follows. Two volumes of RNAprotect (Qiagen) was added to the cultures, and bacterial cells were collected by centrifugation at 5,000 × g for 25 min at 4°C. The dry cell pellet was stored at −80°C overnight. Total RNA was extracted using the RNeasy minikit (Qiagen) with an additional mechanical lysis step using lysing matrix B tubes (MP Biomedicals). RNA samples were treated with Turbo DNase (Ambion) to eliminate contaminating genomic DNA and quantified using a NanoDrop instrument (Thermo Scientific). One microgram of RNA was converted to cDNA using the SuperScript VILO cDNA synthesis kit (Invitrogen). Resulting cDNA libraries were used for downstream analyses. RNA sequencing samples were taken from bacterial cultures grown in THY to late-exponential growth phase (OD600 of ∼0.7-0.8). Erythromycin was used at a concentration of 2 µg/ml. Mitomycin C was added to early-exponential cultures (OD600 of 0.25) at a concentration of 0.2 µg/ml. RNA-sequencing was performed from Ribo-zero (rRNA depleted) Illumina libraries on a single Illumina HiSeq 2500 lane using v4 chemistry from 75 base pair paired-end reads. Reads were mapped to the HKU16 reference genome (alternatively termed HKU QMH11M0907901, GenBank accession number NZ_AFRY01000001) with BWA MEM (version 0.7.16). Relative read counts (per gene) and differential gene expression was determined using DESeq2^59^ (v. 1.26.0) in R (v. 3.6.0). Genes with less than 10 reads across all conditions and samples were removed. P-values were calculated using Wald test and adjusted for multiple testing using Benjamini-Hichberg/false discovery rate. Read counts were visualized using the Integrative Genomics Viewer (IGV) and volcano plots were constructed using ggplot2 (v.3.2.1). To quantify gene expression, total RNA was isolated from bacterial cells harvested at late-exponential growth phase (OD600 of 0.4) in CDM grown in the presence or absence of 2 mM redox-active compounds (L-Cysteine (Cys), dithiothreitol (DTT), reduced glutathione (GSH), oxidized glutathione (GSSG), and tris(2-carboxyethyl)phosphine (TCEP). Reverse transcription-PCR (RT-PCR) was performed using the primers specified in Table S1, using SYBR green master mix (Applied Biosystems) according to the manufacturer’s instructions. All data were analyzed using ViiA7 software (Applied Biosystems). Relative gene expression was calculated using the threshold cycle (2^−ΔΔCT^) method with *gyrA* as the reference housekeeping gene^60^. All reactions were performed in triplicate from 3 independently isolated RNA samples.

### Purification of streptococcal antigens and polyclonal antiserum production

The gene encoding for the DNase Spd1, including nucleotides encoding the predicted signal peptide, was PCR amplified from genomic DNA of HKU16 and cloned into *Nde*I and *Hin*dIII sites of pET-28a. Point mutation of the active site residue Asn145 (Asn145Ala)^61^ was introduced using the QuikChange II site-directed mutagenesis kit (Agilent) to inactivate the Spd1 DNase (see Supplementary Table S2 for primer sequences). WT Spd1 and inactivated Spd1 were produced by 0.5 mM isopropyl β-D-1-thiogalactopyranoside (IPTG)-induced expression in *E. coli* BL21(DE3), purified via nickel affinity chromatography, and His_6_ tags cleaved with His_6_-tagged tobacco etch virus (TEV) protease. The expression plasmids for WT SLO (pET-15b-SLO)^43^ and inactivated SLO carrying P427L and W535A mutations (pET-15b-SLOmut)^62^ were used to produce recombinant protein in *E. coli* BL21(DE3) following the same procedure as for Spd1. Recombinant proteins were analyzed for purity on 12% separating sodium dodecyl sulfate polyacrylamide gel electrophoresis (SDS-PAGE). Inactivated Spd1 and SLO were used to raise antisera in mice. Briefly, 4 - 6 weeks old BALB/c mice (n = 10) were immunized subcutaneously on days 0, 14, 21, and 28 with 30 µg of total protein adjuvanted with alum (Alhydrogel [2%]; Brenntag) at a 1:1 ratio. One week following the last injection, mice were sacrificed and serum was collected for antibody titer analysis using ELISA.

### Detection of exotoxins in GAS supernatants

Bacteria were routinely grown to late-exponential growth phase in CDM or THY where indicated. Filter-sterilized culture supernatants were precipitated with 10% trichloroacetic acid (TCA). TCA precipitates were resuspended in loading buffer (normalized to OD600) in the presence or absence of 100 mM DTT. Samples were boiled for 10 min, subjected to SDS-PAGE, and then transferred to polyvinylidene difluoride (PVDF) membranes for detection of immuno-reactive bands using a LI-COR Odyssey Imaging System (LI-COR Biosciences). The primary antibodies used for the detection of SpeC and SSA protein in GAS culture supernatants were rabbit antibody to SpeC (PCI333, Toxin Technology; 1:1,000 dilution) and affinity-purified rabbit antibody to SSA (produced by Mimotopes; 1:500 dilution)^15^. The murine primary antibody dilutions used for the detection of Spd1 and SLO were 1:1,000 and 1:2,000, respectively. Anti-rabbit IgG (H+L) (DyLight™ 800 4X PEG Conjugate, NEB) or anti-mouse IgG (H+L) (DyLight™ 800 4X PEG Conjugate, NEB) were used as the secondary antibodies.

### Recombinant superantigen purification

The SSA gene, lacking nucleotides encoding the predicted signal peptide, was PCR amplified from the *S. pyogenes* HKU16 chromosome using primers listed in Table S1 and cloned into the *Nco*I and *Bam*HI sites of a modified pET-41a protein expression vector that encodes an engineered tobacco etch virus (TEV) protease site to remove purification tags^63^. The C26S mutation was introduced into the *ssa* gene as above using primers listed in Table S1. Cloning of SpeC into the pET-41a vector was carried out as previously described^20^. Expression of the recombinant SSA and SpeC proteins was induced with 0.2 mM IPTG in *E. coli* BL21(DE3) and purified as described above.

### Superantigen activity as assessed by T cell proliferation assay

Human PBMCs isolated from freshly drawn heparinized venous blood from a healthy adult volunteer were resuspended in complete RPMI (cRPMI; RPMI1640, 10% FBS, 0.1 mM minimal essential media non-essential amino acids, 2 mM L-glutamine, 1 mM sodium pyruvate, 100 U/ml penicillin, 100 ug/ml streptomycin) and seeded at 200,000 cells per well in a 96-well plate. Sterile-filtered GSH dissolved in cRPMI (final concentration of 2 mM), or cRPMI alone, were added to each well 30 minutes prior to the addition of 10-fold serial dilutions of recombinant superantigens. Cells were incubated at 37°C in 5% CO_2_ for 18 h. Spent cell culture supernatant was harvested and analyzed for human IL-2 by ELISA according to the manufacturer’s instructions (eBiosciences).

### *Ex vivo* whole blood model

Freshly drawn heparinized venous blood from a healthy adult volunteer was aliquoted (180 µl) into wells of a 96-well plate. To validate hemolytic activity of SLO, increasing concentrations of recombinant WT SLO were added to give a final volume of 200 µl per well and incubated at 37°C for 2 h with 5% CO_2_. For bacterial infections, GAS strains were grown to late-exponential growth phase in CDM (OD600 of 0.4), resuspended in Hanks Balanced Salt Solution (HBSS) at ∼ 1 × 10^8^ CFU/ml, and then added to whole blood to give a final volume of 200 µl (∼ 2 × 10^6^ CFU). Growth of GAS strains was assessed 2 h post-infection by plating serial dilutions for enumeration of CFUs. Plasma samples for detection of hemolysis and glutathione release were obtained 4 h post-infection by centrifugation at 4,800 × g for 15 min at 4°C. Controls included for each experiment were whole blood treated with HBSS (mock), or blood lyzed with 0.1% Triton X-100.

### Co-culture of *S. pyogenes* with human pharyngeal cells

Human nasopharyngeal carcinoma epithelial cells Detroit 562 (D562; ATCC CCL-138, Lot 70004014) were cultured at 37°C under a 5% CO_2_/20% O_2_ atmosphere in EMEM supplemented with 10% FBS in tissue culture vessels (Greiner Bio-one). At 90% confluency, cells were trypsinized and handled according to manufacturer’s instructions. D562 cells were utilized for experiments at passage 8 and seeded at a density of ∼ 1.2 × 10^5^ viable cells per well in 24-well tissue culture plates, or ∼ 2.5 × 10^5^ viable cells per well in 12-well plates 24 h prior to infection to allow the formation of confluent monolayers. Cells were grown at 37°C under 5% CO_2_ until they formed a confluent monolayer. Immediately prior to infection, the cell culture medium was removed, and replaced with fresh medium. Increasing concentrations of recombinant WT SLO were added to cell monolayers and incubated for 2 h. GAS strains were grown to late-exponential growth phase in CDM (OD600 of 0.4), resuspended in cell culture medium, and then added to cell monolayers at a multiplicity of infection of 20. Controls included for each experiment were cells not exposed to bacteria or SLO (mock), or cells lyzed with 0.2% Triton X-100. At 2 h post-infection, plates were centrifuged at 500 × g for 5 min, then media was aspirated and stored at −80°C until further processing.

### Quantitative assessment of cell death and glutathione release

SLO-induced hemolysis in whole blood was determined after collecting plasma samples and diluting 1:10 in PBS. The amount of hemoglobin was measured spectrophotometrically at 405 nm. D562 membrane disruption was quantified by measuring lactate dehydrogenase (LDH) release from cell supernatants, using CytoTox96 Non-Radioactive Cytotoxicity Assay (Promega; G1781), as per the manufacturer’s instructions. Sample absorbance was measured spectrophotometrically at 490 nm. Glutathione release was measured using the GSH-Glo glutathione assay (Promega; V6912), as per the manufacturer’s instructions, with the modification of mixing undiluted samples 1:1 with 2 mM TCEP in wells of a white 96-well plate (Greiner Bio-one) prior to use. Luminescent intensity of each sample was measured using a FLUOstar Omega Microplate Reader (BMG Labtech). Sample readings were analyzed by Prism 7 software and divided by the positive control for cell lysis to give a percentage of total hemolysis and cell death (LDH) for each sample.

### Neutrophil killing assay

Human neutrophils were isolated from fresh heparinized whole blood using PolymorphPrep density gradient centrifugation (Axis-Shield) as per manufacturer’s instructions. Following neutrophil harvest, hypotonic lysis was performed to remove residual erythrocytes. Purified neutrophils were infected with GAS at a multiplicity of infection of 10 (1 × 10^6^ cells/ml neutrophils:1 × 10^5^ bacterial CFU/ml), centrifuged for 5 min at 370 × g to synchronize phagocytosis, and then incubated for 30 min at 37°C under 5% CO_2_. Control wells contained bacteria only. Infected neutrophils were then lysed and serially diluted in sterile Milli-Q water, then plated on THY agar. Following overnight incubation at 37°C, bacterial survival was calculated as the average total number of CFUs following incubation in the presence of neutrophils divided by CFUs in control wells.

### NETs degradation assay

Freshly isolated purified human neutrophils were seeded on 12 mm Poly-D-lysine-coated (0.01 % solution overnight; Sigma-Aldrich; P7405) coverslips at a concentration of 1 × 10^6^ cells/ml (5 × 10^5^ cells/mL per well) in a 24-well tissue culture plate. Neutrophils were stimulated with 25 nM phorbol 12-myristate 13-acetate (PMA) (Cayman Chemical; 10008014), centrifuged for 5 min at 370 × g, and incubated for 3 h at 37°C under 5% CO_2_ to induce NET formation. Cell culture media was then removed, and the PMA-stimulated neutrophils were infected with fluorescent GAS strains diluted in RPMI media containing 2% heat inactivated autologous human plasma and 5 mM MgCl_2_ at a multiplicity of infection of 10 (1 × 10^7^ bacterial CFU/ml: 1 × 10^6^ cells/ml PMN). Infected plates were centrifuged at 370 × g for 5 min to promote cell interaction and then incubated for an additional 30 min at 37°C under 5% CO_2_. Bovine pancreas DNase I at 5 µg/ml (Sigma; D5025) was used as a positive control to confirm NET degradation, while wells containing medium was used to confirm the formation of NETs. Cells were washed once with PBS, followed by fixation with 4% paraformaldehyde for 15 min at room temperature. After two washes, cells were incubated with 1 mM SYTOX Orange Nucleic Acid Stain (Molecular Probes; S11368) for 15 min in the dark at room temperature to stain for NETs. After washing in 5% (v/v) PBS, coverslips were embedded in Fluorescent Mounting medium (Dako; S30230) on microscopic glass slides and dried overnight in the dark at room temperature. Slides were stored at 4°C until images were acquired. Samples were recorded using a Leica TCS SP8 Lightning confocal laser scanning microscope (Leica Microsystems) with a 63 × oil immersion objective. GFP and SYTOX Orange were excited with 488 and 561 nm lasers, respectively, with images captured using sequential scanning. For each sample, a minimum of five randomly selected images per independent experiment performed in duplicate were acquired. For figure production, images were processed using ImageJ software (version 1.8.0) and the Enhance Local Contrast function was used to improve images for better visualization. For quantification of NET DNA degradation, the cell imaging analysis software CellProfiler (version 3.1.9) was employed. The percentage area of NETs per image was calculated as the area of neutrophil nuclei subtracted from the total area stained with SYTOX Orange. Images used for NET quantification were unenhanced.

### HLA-B6 murine nasopharyngeal colonization model

For nasopharyngeal infection^20,21^, sex- and age-matched (9- to 13-week-old) transgenic C57BL/6J mice expressing human major histocompatibility complex II molecules DR4/DQ8 and human CD4 (HLA-B6)^42^ were infected with ∼ 1 × 10^8^ CFU per 15 µl using 7.5 µl to inoculate each nostril under methoxyflurane inhalation anesthetic. For infection, bacteria were cultured to late-exponential growth phase (OD600 of 0.4) in CDM supplemented with 2 mM of L-Cys, washed and concentrated in CDM. Sham-treated mice only received CDM. Mice were sacrificed 48 h post-infection, and the combined nasal turbinates, including the nasal associated lymphoid tissue and nasal turbinates, were removed. Tissue was homogenized in HBSS in lysing matrix F tubes (MP Biomedicals), serially diluted, and plated on horse blood agar for enumeration of beta-hemolytic CFUs.

### Statistical analysis

All statistical analysis was completed using Prism software (GraphPad). Significance was calculated using, where indicated, the Student’s t test, one-way ANOVA with Dunnett’s multiple comparisons post-hoc test, and the Kruskal-Wallis test with the Dunn’s multiple comparisons post-hoc test. A p value less than 0.05 was determined to be statistically significant.

### Accession codes

Illumina read data are available on NCBI under the sample accession numbers relating to the three conditions (in triplicate): THY (ERS1091539, ERS1091548, ERS1091557); THY plus erythromycin (ERS1091542, ERS1091551, ERS1091560); THY plus mitomycin C (ERS1091545, ERS1091554, ERS1091563).

### Ethics statement

The human ethics protocol for the isolation of human blood from healthy volunteers for use in T cell activation assays was approved by the Health Sciences Research Ethics Board at Western University (Ontario, Canada) (Protocol #110859). Human blood donation for use in whole blood proliferation assays, neutrophil killing assays and NET degradation assays were conducted in accordance with the Australian National statement on ethical conduct in human research^64^, in compliance with the regulations governing experimentation on humans, and was approved by the University of Queensland medical research ethics committee (2010001586) and the University of Wollongong Human Research Ethics Committee (HE08/250). Animal experiments were performed according to the Australian code of practice for the care and use of animals for scientific purposes. Permission was obtained from the University of Queensland ethics committee to undertake this work (SCMB/140/16/NHMRC).

## References

1 Katz, A. R. & Morens, D. M. Severe streptococcal infections in historical perspective. Clin Infect Dis 14, 298–307 (1992)

2 Morens, D. M., Folkers, G. K. & Fauci, A. S. The challenge of emerging and re-emerging infectious diseases. Nature 430, 242–249 (2004)

3 Luk, E. Y. et al. Scarlet fever epidemic, Hong Kong, 2011. Emerg Infect Dis 18, 1658–1661 (2012)

4 Tse, H. et al. Molecular characterization of the 2011 Hong Kong scarlet fever outbreak. J Infect Dis 206, 341–351 (2012)

5 Turner, C. E. et al. Scarlet fever upsurge in England and molecular-genetic analysis in North-West London, 2014. Emerg Infect Dis 22, 1075–1078 (2016)

6 Chalker, V. et al. Genome analysis following a national increase in scarlet fever in England 2014. BMC Genomics 18, 224 (2017)

7 Park, D. W. et al. Incidence and characteristics of scarlet fever, South Korea, 2008-2015. Emerg Infect Dis 23, 658–661 (2017)

8 Liu, Y. et al. Resurgence of scarlet fever in China: a 13-year population-based surveillance study. Lancet Infect Dis 18, 903–912 (2018)

9 Lamagni, T. et al. Resurgence of scarlet fever in England, 2014-16: a population-based surveillance study. Lancet Infect Dis 18, 180–187 (2018)

10 Lynskey, N. N. et al. Emergence of dominant toxigenic M1T1 *Streptococcus pyogenes* clone during increased scarlet fever activity in England: a population-based molecular epidemiological study. Lancet Infect Dis 19, 1209–1218 (2019)

11 Yung, C. F. & Thoon, K. C. A 12 year outbreak of scarlet fever in Singapore. Lancet Infect Dis 18, 942 (2018)

12 Walker, M. J. et al. Detection of epidemic scarlet fever group A *Streptococcus* in Australia. Clin Infect Dis 69, 1232–1234 (2019)

13 Demczuk, W., Martin, I., Domingo, F. R., MacDonald, D. & Mulvey, M. R. Identification of *Streptococcus pyogenes* M1UK clone in Canada. Lancet Infect Dis 19, 1284–1285 (2019)

14 Walker, M. J. & Brouwer, S. Scarlet fever makes a comeback. Lancet Infect Dis 18, 128–129 (2018)

15 Davies, M. R. et al. Emergence of scarlet fever *Streptococcus pyogenes emm12* clones in Hong Kong is associated with toxin acquisition and multidrug resistance. Nat Genet 47, 84–87 (2015)

16 Ben Zakour, N. L. et al. Transfer of scarlet fever-associated elements into the group A *Streptococcus* M1T1 clone. Sci Rep 5, 15877 (2015)

17 You, Y. et al. Scarlet fever epidemic in China caused by *Streptococcus pyogenes* serotype M12: epidemiologic and molecular analysis. EBioMedicine 28, 128–135 (2018)

18 Silva-Costa, C., Carrico, J. A., Ramirez, M. & Melo-Cristino, J. Scarlet fever is caused by a limited number of *Streptococcus pyogenes* lineages and is associated with the exotoxin genes *ssa, speA* and *speC*. Pediatr Infect Dis J 33, 306–310 (2014)

19 Proft, T. a. F. J.D. Streptococcal superantigens: biological properties and potential role in disease. In: Ferretti, JJ, Stevens, DL, Fischetti, VA, eds. Streptococcus pyogenes: Basic Biology to Clinical Manifestations [Internet]. Oklahoma City:University of Oklahoma Health Sciences Center, 445–486 (2016)

20 Kasper, K. J. et al. Bacterial superantigens promote acute nasopharyngeal infection by *Streptococcus pyogenes* in a human MHC Class II-dependent manner. PLoS Pathog 10, e1004155–e1004155 (2014)

21 Zeppa, J. J. et al. Nasopharyngeal infection by *Streptococcus pyogenes* requires superantigen-responsive Vbeta-specific T cells. Proc Natl Acad Sci U S A 114, 10226–10231 (2017)

22 Fossum, G. H., Lindbaek, M., Gjelstad, S., Dalen, I. & Kvaerner, K. J. Are children carrying the burden of broad-spectrum antibiotics in general practice? Prescription pattern for paediatric outpatients with respiratory tract infections in Norway. BMJ Open 3, pii: e002285 (2013)

23 Bisno, A. L. Acute pharyngitis. N Engl J Med 344, 205–211 (2001)

24 Banks, D. J., Lei, B. & Musser, J. M. Prophage induction and expression of prophage-encoded virulence factors in group A *Streptococcus* serotype M3 strain MGAS315. Infect Immun 71, 7079–7086 (2003)

25 Broudy, T. B., Pancholi, V. & Fischetti, V. A. Induction of lysogenic bacteriophage and phage-associated toxin from group A streptococci during coculture with human pharyngeal cells. Infect Immun 69, 1440–1443 (2001)

26 Broudy, T. B., Pancholi, V. & Fischetti, V. A. The *in vitro* interaction of *Streptococcus pyogenes* with human pharyngeal cells induces a phage-encoded extracellular DNase. Infect Immun 70, 2805–2811 (2002)

27 Sundberg, E. J. et al. Structures of two streptococcal superantigens bound to TCR beta chains reveal diversity in the architecture of T cell signaling complexes. Structure 10, 687–699 (2002)

28 De Marzi, M. C. et al. Cloning, expression and interaction of human T-cell receptors with the bacterial superantigen SSA. Eur J Biochem 271, 4075–4083 (2004)

29 Proft, T., Moffatt, S. L., Berkahn, C. J. & Fraser, J. D. Identification and characterization of novel superantigens from *Streptococcus pyogenes*. J Exp Med 189, 89–102 (1999)

30 Tweten, R. K. Cholesterol-dependent cytolysins, a family of versatile pore-forming toxins. Infect Immun 73, 6199–6209 (2005)

31 Schafer, F. Q. & Buettner, G. R. Redox environment of the cell as viewed through the redox state of the glutathione disulfide/glutathione couple. Free Radic Biol Med 30, 1191–1212 (2001)

32 Alouf, J. E. Streptococcal toxins (streptolysin O, streptolysin S, erythrogenic toxin). Pharmacol Ther 11, 661–717 (1980)

33 Bessen, D. E. Tissue tropisms in group A *Streptococcus*: what virulence factors distinguish pharyngitis from impetigo strains? Curr Opin Infect Dis 29, 295–303 (2016)

34 Aziz, R. K. et al. Mosaic prophages with horizontally acquired genes account for the emergence and diversification of the globally disseminated M1T1 clone of *Streptococcus pyogenes*. J Bacteriol 187, 3311–3318 (2005)

35 Sumby, P. et al. Evolutionary origin and emergence of a highly successful clone of serotype M1 group A *Streptococcus* involved multiple horizontal gene transfer events. J Infect Dis 192, 771–782 (2005)

36 Walker, M. J. et al. DNase Sda1 provides selection pressure for a switch to invasive group A streptococcal infection. Nat Med 13, 981–985 (2007)

37 Afshar, B. et al. Enhanced nasopharyngeal infection and shedding associated with an epidemic lineage of *emm3* group A *Streptococcus*. Virulence 8, 1390–1400 (2017)

38 Brinkmann, V. et al. Neutrophil extracellular traps kill bacteria. Science 303, 1532–1535 (2004)

39 Buchanan, J. T. et al. DNase expression allows the pathogen group A *Streptococcus* to escape killing in neutrophil extracellular traps. Curr Biol 16, 396–400 (2006)

40 Park, H. S. et al. Primary induction of CD4 T cell responses in nasal associated lymphoid tissue during group A streptococcal infection. Eur J Immunol 34, 2843–2853 (2004)

41 Wang, B. et al. Induction of TGF-beta1 and TGF-beta1-dependent predominant Th17 differentiation by group A streptococcal infection. Proc Natl Acad Sci U S A 107, 5937–5942 (2010)

42 Chen, Z. et al. Humanized transgenic mice expressing HLA DR4-DQ3 haplotype: reconstitution of phenotype and HLA-restricted T-cell responses. Tissue Antigens 68, 210–219 (2006)

43 Timmer, A. M. et al. Streptolysin O promotes group A *Streptococcus* immune evasion by accelerated macrophage apoptosis. J Biol Chem 284, 862–871 (2009)

44 Uchiyama, S. et al. Streptolysin O rapidly impairs neutrophil oxidative burst and antibacterial responses to group A *Streptococcus*. Front Immunol 6, 581 (2015)

45 Zhu, L. et al. Contribution of secreted NADase and streptolysin O to the pathogenesis of epidemic serotype M1 *Streptococcus pyogenes* infections. Am J Pathol 187, 605–613 (2017)

46 Hsieh, Y. C. & Huang, Y. C. Scarlet fever outbreak in Hong Kong, 2011. J Microbiol Immunol Infect 44, 409–411 (2011)

47 Chen, M. et al. Outbreak of scarlet fever associated with *emm12* type group A *Streptococcus* in 2011 in Shanghai, China. Pediatr Infect Dis J 31, e158–162 (2012)

48 Lau, E. H., Nishiura, H., Cowling, B. J., Ip, D. K. & Wu, J. T. Scarlet fever outbreak, Hong Kong, 2011. Emerg Infect Dis 18, 1700–1702 (2012)

49 Yang, P. et al. Characteristics of group A *Streptococcus* strains circulating during scarlet fever epidemic, Beijing, China, 2011. Emerg Infect Dis 19, 909–915 (2013)

50 Brouwer, S., Lacey, J. A., You, Y., Davies, M. R. & Walker, M. J. Scarlet fever changes its spots. Lancet Infect Dis 19, 1154–1155 (2019)

51 Brosnahan, A. J., Mantz, M. J., Squier, C. A., Peterson, M. L. & Schlievert, P. M. Cytolysins augment superantigen penetration of stratified mucosa. J Immunol 182, 2364–2373 (2009)

52 Veeravalli, K., Boyd, D., Iverson, B. L., Beckwith, J. & Georgiou, G. Laboratory evolution of glutathione biosynthesis reveals natural compensatory pathways. Nat Chem Biol 7, 101–105 (2011)

53 Brenot, A., King, K. Y., Janowiak, B., Griffith, O. & Caparon, M. G. Contribution of glutathione peroxidase to the virulence of *Streptococcus pyogenes*. Infect Immun 72, 408–413 (2004)

54 Reniere, M. L. Reduce, induce, thrive: bacterial redox sensing during pathogenesis. J Bacteriol 200 pii: e00128–18 (2018)

55 Ku, J. W. & Gan, Y. H. Modulation of bacterial virulence and fitness by host glutathione. Curr Opin Microbiol 47, 8–13 (2018)

56 Commons, R. et al. Superantigen genes in group A streptococcal isolates and their relationship with *emm* types. J Med Microbiol 57, 1238–1246 (2008)

57 Chochua, S. et al. Emergent invasive group A *Streptococcus dysgalactiae* subsp. *equisimilis*, United States, 2015-2018. Emerg Infect Dis 25, 1543–1547 (2019)

58 Barnett, T. C., Daw, J. N., Walker, M. J. & Brouwer, S. Genetic manipulation of group A *Streptococcus* – gene deletion by allelic replacement. In: T. Proft and J. Loh (Ed) Group A Streptococcus: Methods and Protocols. Springer, Heidelberg, in press (2020)

59 Love, M. I., Huber, W. & Anders, S. Moderated estimation of fold change and dispersion for RNA-seq data with DESeq2. Genome Biol 15, 550 (2014)

60 Brouwer, S. et al. Endopeptidase PepO regulates the SpeB cysteine protease and is essential for the virulence of invasive M1T1 *Streptococcus pyogenes*. J Bacteriol 200, pii: e00654–17 (2018)

61 Korczynska, J. E., Turkenburg, J. P. & Taylor, E. J. The structural characterization of a prophage-encoded extracellular DNase from *Streptococcus pyogenes*. Nucleic Acids Res 40, 928–938 (2012)

62 Rivera-Hernandez, T. et al. Differing efficacies of lead group A streptococcal vaccine candidates and full-length M protein in cutaneous and invasive disease models. mBio 7, pii: e00618–16 (2016)

63 Rahman, A. K. M. N. U. et al. Molecular basis of TCR selectivity, cross-reactivity, and allelic discrimination by a bacterial superantigen: integrative functional and energetic mapping of the SpeC-V beta 2.1 molecular interface. J Immunol 177, 8595–8603 (2006)

64 National statement on ethical conduct in human research. National Health and Medical Research Council, Canberra, Australia, https://www.nhmrc.gov.au/guidelines-publications/e72 (2015).

65 Palmer, M. The family of thiol-activated, cholesterol-binding cytolysins. Toxicon 39, 1681–1689 (2001)

66 Cywes Bentley, C., Hakansson, A., Christianson, J. & Wessels, M. R. Extracellular group A *Streptococcus* induces keratinocyte apoptosis by dysregulating calcium signalling. Cell Microbiol 7, 945–955 (2005)

