## Supplementary Information for "Prophage exotoxins enhance colonization fitness in epidemic scarlet fever-causing *Streptococcus pyogenes*"

**Construction of reporter strains**

The plasmid-based reporter system (pLZ12Km2-P23R:TA, Addgene plasmid gift from Associate Professor Thomas Proft, University of Auckland, New Zealand) described in[1] was used to construct plasmid pLZ12Km2-P23R:TA:GFP. Maintenance plasmid pUC57-RBSGFP containing the ribosomal binding site (RBS) and *gfp* gene from pDCerm-GFP[2] was synthesized commercially by Genscript. pLZ12Km2-P23R:TA was digested with *Not*I, and pUC57-RBSGFP was incubated with *Not*I to excise the RBS and *gfp* gene from pUC57-RBSGFP. The excised RBS and *gf*p were then ligated into digested pLZ12km2-P23R:TA to generate pLZ12Km2-P23R:TA:GFP, which was used for transformation of electrocompetent HKU16 cells.

### Supplementary Figures

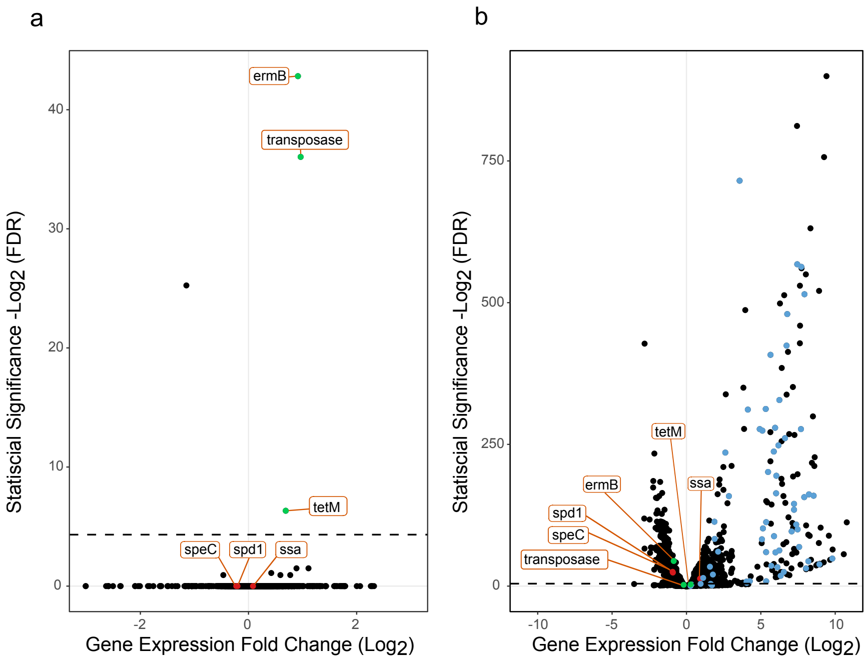


**Fig. S1:** RNA-seq transcriptomes of *S. pyogenes emm*12 strain HKU16 grown in THY medium compared with (**a**) THY supplemented with 2 µg/ml erythromycin and (**b**) THY supplemented with 0.2 µg/ml mitomycin C. Volcano plots represent differentially expressed genes of erythromycin or mitomycin C supplemented cultures relative to cultures grown in THY medium alone. Each dot represents a gene expression fold change (horizontal axis) with respect to statistical significance (vertical axis). Genes relating to prophage ϕHKU.vir virulence factors (red dots) and ICE-HKU*emm*12 resistance genes (green dots) are annotated as indicated using orange boxes. Genes corresponding to prophage ΦHKU.vir genes are colored blue in (**b**). Dashed line indicates a false discovery rate (FDR) of -Log$_2$ 0.05.

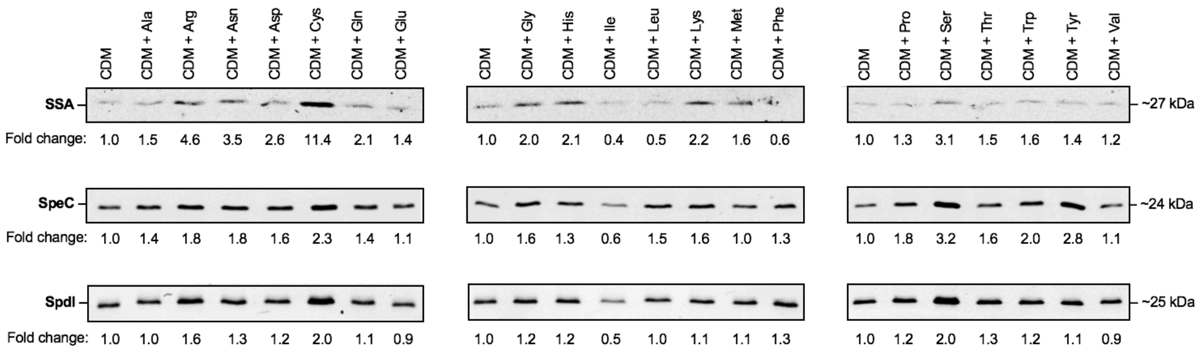


**Fig. S2:** Small molecule screen of all 20 amino acids used to identify cysteine as a factor specifically enhancing release of the exotoxin SSA by HKU16 grown in chemically defined medium (CDM).

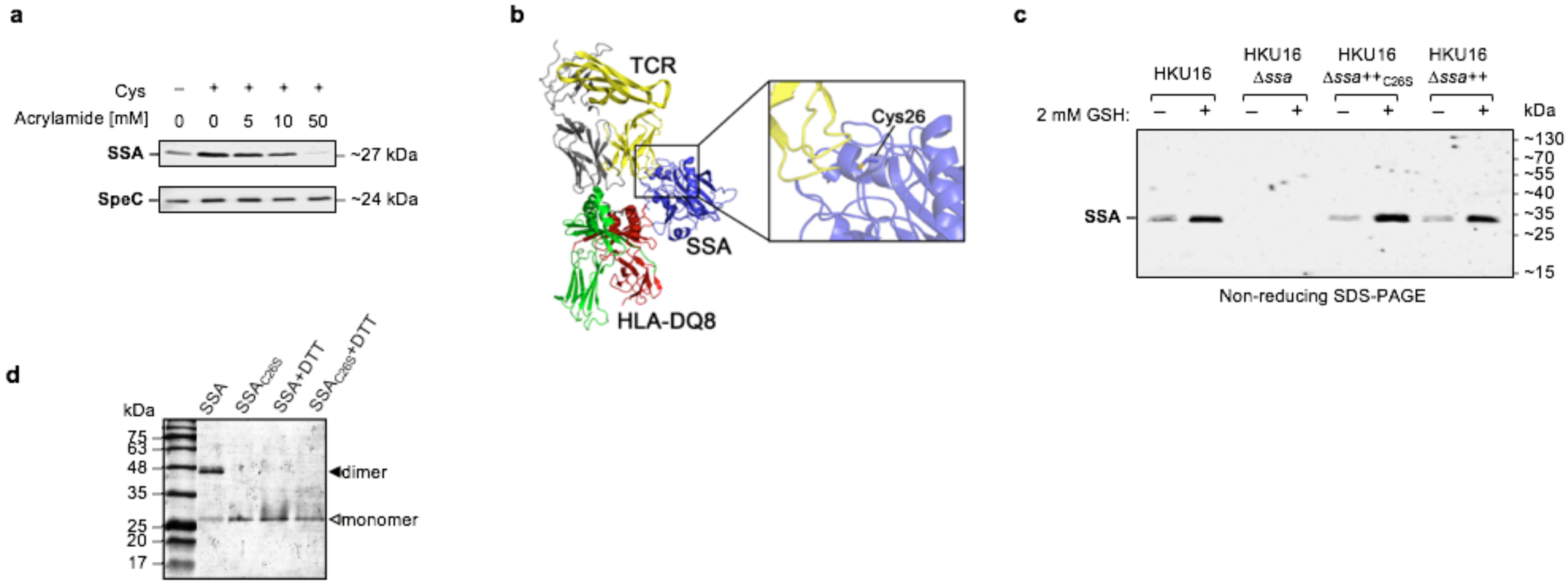


**Fig. S3:** (**a**) Western immunoblot detection of secreted SSA and SpeC after supplementation of CDM with 2 mM Cys pre-treated with increasing concentrations of acrylamide. (**b**) Ribbon diagram representation of the modelled SSA-mediated T cell activation complex. The model was generated by superposition of the SSA (PDB 1BXT)[3] and HLA-DQ8 (PDB 1JK8)[4] crystal structures onto the co-crystal of SpeA in complex with TCRβ (PDB 1L0Y) and the co-crystal of SEB in complex with HLA-DR4 (PDB 1SEB)[5]. Colors are as follows: SSA, *blue*; TCR β-chain *yellow*; TCR α-chain *grey*; MHC α-chain, *red*; MHC β-chain, *green*. The inset highlights the location of the free Cys26 within the TCR-SSA interface. The image was generated using the PyMOL Molecular Graphics System, version 1.3 Schrödinger (www.pymol.org/). (**c**) Immunoblot detection of SSA secreted by indicated HKU16 strains grown in CDM supplemented with 2 mM of GSH following non-reducing SDS-PAGE. (**d**) PAGE profile (lacking SDS and samples were not boiled) of purified WT SSA and mutant $SSA_{C26S}$ under non-reducing and reducing conditions, respectively.

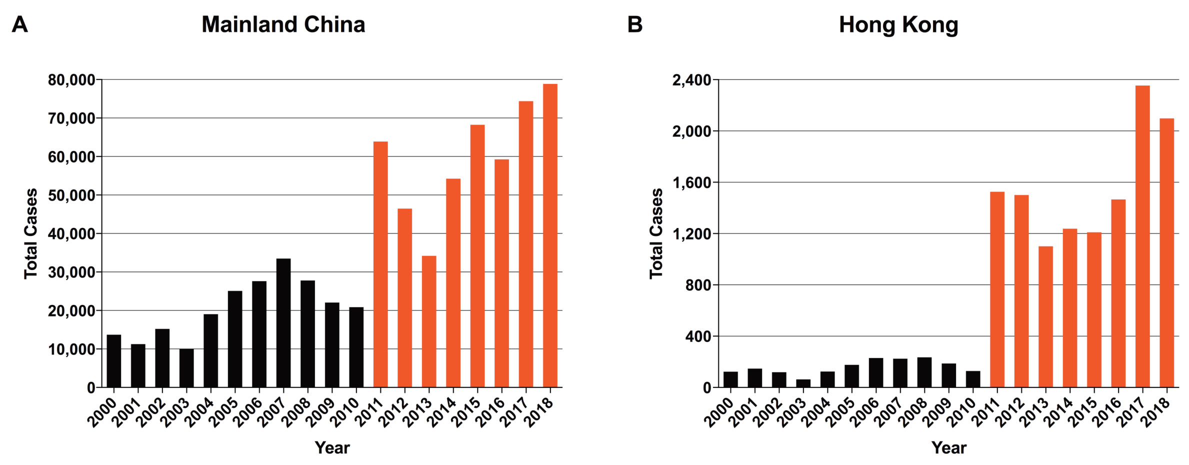


**Fig. S4:** Annual reported scarlet fever cases in (A) Mainland China and (B) Hong Kong. Data were obtained from the National Bureau of Statistics of China (http://data.stats.gov.cn; accessed August 20, 2019), and from the Hong Kong Centre for Health Protection website (http://www.chp.gov.hk/en/notifiable1/10/26/43.html; August 20, 2019), respectively. Ongoing resurgence of scarlet fever is highlighted in red.

**Table S1:** List of bacterial strains, plasmids and primers used in this study.

| Bacterial strains | Description | Reference/Source |
|---|---|---|
| ***E. coli*** | | |
| MC1061 | Laboratory cloning strain | [6] |
| XL1-blue | Laboratory cloning strain | Stratagene |
| BL21(DE3) | Laboratory expression strain | Stratagene |
| ***S. pyogenes*** | | |
| HKU16 | Hong Kong *S. pyogenes emm12* scarlet fever isolate | [7] |
| HKU16Δ*ssa* | HKU16Δ*ssa* isogenic mutant strain | This study |
| HKU16Δ*ssa*++ | HKU16Δ*ssa*::*ssa*-complemented strain | This study |
| HKU16Δ*ssa*(C26S) | HKU16Δ*ssa*::*ssa*-complemented strain containing a C26S substitution | This study |
| HKU16Δ*speC* | HKU16Δ*speC* isogenic mutant strain | This study |
| HKU16Δ*spdI* | HKU16Δ*spdI* isogenic mutant strain | This study |
| HKU16Δ*spdI*++ | HKU16Δ*spdI*::*spdI*-complemented strain | This study |
| HKU16Δ*ssa*/*speC* | HKU16Δ*ssa*/*speC* double isogenic mutant strain | This study |
| HKU16Δ*ssa*/*spdI* | HKU16Δ*ssa*/*spdI* double isogenic mutant strain | This study |
| HKU16Δ*speC*/*spdI* | HKU16Δ*speC*/*spdI* double isogenic mutant strain | This study |
| HKU16Δ*ssa*/*speC*/*spdI* | HKU16Δ*ssa*/*speC*/*spdI* triple isogenic mutant strain | This study |
| HKU16Δ*ssa*/*speC*/*spdI*++ | HKU16Δ*ssa*/*speC*/*spdI*::*ssa*/*speC*/*spdI* complemented strain | This study |
| HKU16Δ*slo* | HKU16Δ*slo* isogenic mutant strain | This study |
| HKU16-GFP | HKU16 carrying *gfp* reporter pLZ12Km2-P23R:TA:GFP | This study |
| HKU16Δ*spdI*-GFP | HKU16Δ*spdI* carrying *gfp* reporter pLZ12Km2-P23R:TA:GFP | This study |
| HKU16Δ*spdI*++-GFP | HKU16Δ*spdI*::*spdI*-complemented strain carrying *gfp* reporter pLZ12Km2-P23R:TA:GFP | This study |
| **Plasmids** | | |
| ***Expression plasmids*** | | |
| pET-28a | Expression plasmid::kanamycin$^R$ | Novagen |
| pET-28a-Spd1 | pET-28a+SpdI expression construct | This study |
| pET-28a-Spd1_N145A | pET-28a+SpdI expression construct containing a N145A substitution | This study |
| pET-151 | Directional TOPO expression plasmid, ampicillin$^R$ | Invitrogen |
| pET-151-SSA_N20D/N23A/Y89A/Y94A | pET-151+SSA expression construct containing N20D/N23A/Y89A/Y94A substitutions | This study |
| pET-15b-SLO | pET-15b+SLO expression construct | [8] |
| pET-15b-SLOmut | pET-15b+SLO expression construct containing P427L/W535A substitutions | [9] |
| pET-41a | Expression plasmid:: kanamycin$^R$ | Novagen |
| pET-41a-SSA | pET-41a+SSA expression construct | This study |
| pET-41a-SSA_C26S | pET-41a+SSA expression construct containing a C26S substitution | This study |
| pET-41a-SpeC | pET-41a+SpeC expression construct | [10] |
| ***Mutagenesis plasmids*** | | |

| | | |
|---|---|---|
| pLZts | Temperature-sensitive shuttle plasmid, spectinomycin$^{R}$ | 11 |
| pLZts-*ssa*_KO | pLZts+*ssa* knockout construct | This study |
| pLZts-*ssa*_complemented | pLZts+*ssa* complementation construct | This study |
| pLZts-*ssa*_complemented_C26S | pLZts+*ssa* complementation construct containing a C26S substitution | This study |
| pLZts-*speC*_KO | pLZts+*speC* knockout construct | This study |
| pLZts-*spdI*_KO | pLZts+*spdI* knockout construct | This study |
| pLZts-*spdI*_complemented | pLZts+*spdI* complementation construct | This study |
| pLZts-*ssa*/*speC*_KO | pLZts+*ssa*/*speC* double knockout construct | This study |
| pLZts-*ssa*/*spdI*_KO | pLZts+*ssa*/*spdI* double knockout construct | This study |
| pLZts-*speC*/*spdI*_KO | pLZts+*speC*/*spdI* double knockout construct | This study |
| pLZts-*ssa*/*speC*/*spdI*_KO | pLZts+*ssa*/*speC*/*spdI* triple knockout construct | This study |
| pLZts-*ssa*/*speC*/*spdI*_complemented | pLZts+*ssa*/*speC*/*spdI* complementation construct | This study |
| pLZts-*slo*_KO | pLZts+*slo* knockout construct | This study |
| ***Reporter plasmids*** | | |
| pLZ12Km2-P23R-TA:GFP | Plasmid-based GFP reporter system | This study |
| **Primer Name** | **Sequence (5'–3')** | |
| ***Primers for protein expression constructs*** | | |
| **Spd1** | | |
| NdeI_*spdI*_SS_pET28a_F | taacatatgatgaaattatctaaacaaaaggcaagtttgcttac | |
| HindIIIstop_*spdI*_pET28a_R | ctgaagcttttattagtttttaggagtggcagttccatttaaatag | |
| *spdI*_N145A_F | cctgaataagcacctgtggctagccaggctgtcattgcc | |
| *spdI*_N145A_R | ggcaatgacagcctggctagccacaggtgcttattcagg | |
| **SSA** | | |
| NcoI_pET-41a_*ssa*_F | gcgccatggcaagtagtcagcctgaccctact | |
| BamHI_pET-41a_*ssa*_R | taagggatccttatttttggtaaggtgaac | |
| *ssa*_C26S_t154a_F | ctacaaaatggttatcatataaacttctcaaattacccataacaccagt | |
| *ssa*_C26S_t154a_R | actggtgttatgggtaatttgagaagtttatatgataaccattttgtag | |
| ***Primers for HKU16 gene chromosomal deletion and complementation constructs*** | | |
| ***ssa*** | | |
| *ssa*_KO-S-F | ttggtcgtcagactgatgggcccatgctacaaggggagagaatc | |
| *ssa*_KO-S-R | gaacctctatgagtattcttattcttttattcatttggctacctcttatatatttaaaac | |
| *ssa*_KO-AS-F | aagaatactcatagaggttcaccttaccaaaaaataaaagaaaataac | |
| ssa_KO-AS-R | cataacctgaaggaagatctcttaaataacaaaattattctagaaaaagatatcg | |
| ***speC*** | | |
| *speC*_KO-S-F | ttggtcgtcagactgatgggccccactaaaataaattatgaccctg | |
| *speC*_KO-S-R | tatcgaaatgtttgatgatgttaatctttttcattttttc | |
| *speC*_KO-AS-F | catcatcaaacatttcgatatttatcttgaaaaataattc | |
| *speC*_KO-AS-R | cataacctgaaggaagatctgtttgatattacaactaataaaaaacaag | |

| | |
|---|---|
| ***spdI*** | |
| *spdI*_KO-S-F | ttggtcgtcagactgatgggcccaacaaaaaagaaccttaatatgg |
| *spdI*_KO-S-R | cagttccatttgccttttgtttagataatttc |
| *spdI*_KO-AS-F | acaaaaggcaaatggaactgccactcctaaaaac |
| *spdI*_KO-AS-R | cataacctgaaggaagatctttttaaagtcaatttcctggaaagttac |
| ***slo*** | |
| *slo*_KO-S-F | ttggtcgtcagactgatgggcccggtgaccataaaaaagtaac |
| *slo*_KO-S-R | tcgaaccatattttttgttagacatgtccttc |
| *slo*_KO-AS-F | ctaacaaaaaatatggttcgattacttataagtag |
| *slo*_KO-AS-R | cataacctgaaggaagatctgagatttccagccttcattataac |
| ***ssa/speC***[1] | |
| *ssa_speC*_KO-S-F | ttggtcgtcagactgatgggcccattcataggataatcacactag |
| *ssa_speC*_KO-S-R | taacatcatcaaacatttcgatatttatcttgaaaaataattc |
| *speC*_KO-S-R | tatcgaaatgtttgatgatgttaatctttttcattttttc |
| *ssa_speC*_KO-AS-R | cataacctgaaggaagatctctaccttttcacatatccaac |
| ***ssa/spdI***[1] | |
| *ssa_spdI*_KO-S-F | ttggtcgtcagactgatgggccctaaagtcaatttcctggaaagttac |
| *ssa_spdI*_KO-S-R | acaaaaggcaggaactgccactcctaaaaac |
| *ssa_spdI*_KO-AS-F | tggcagttcctgccttttgtttagataatttc |
| *ssa_spdI*_KO-AS-R | cataacctgaaggaagatctatcatttgctatcattgcc |
| ***speC/spdI*** | |
| *speC_spdI*_KO-S-F | ttggtcgtcagactgatgggccctcaataattctccgtacgag |
| *speC_spdI*_KO-S-R | acaaaaggcacatttcgatatttatcttgaaaaataattc |
| *speC_spdI*_KO-AS-F | tatcgaaatgtgccttttgtttagataatttc |
| *ssa_spdI*_KO-AS-R | cataacctgaaggaagatctatcatttgctatcattgcc |
| ***ssa/speC/spdI***[1] | |
| *ssa_speC*_KO-S-F | ttggtcgtcagactgatgggcccattcataggataatcacactag |
| *speC_spdI*_KO-S-R | acaaaaggcacatttcgatatttatcttgaaaaataattc |
| *speC_spdI*_KO-AS-F | tatcgaaatgtgccttttgtttagataatttc |
| *ssa_spdI*_KO-AS-R | cataacctgaaggaagatctatcatttgctatcattgcc |
| ***Primers for quantitative real time PCR*** | |
| qRTPCR-*gyrA*-F | cgacttgtctgaacgccaaa |
| qRTPCR-*gyrA*-R | gtcagcaatcaaggccaaca |
| qRTPCR-*ssa*-F | gcctgaccctactccagaac |
| qRTPCR-*ssa*-R | agctgacctgtggatcttaca |
| qRTPCR-*speC*-F | ccgaaatgtcttatgaggcctc |
| qRTPCR-*speC*-R | agcaggcgtaattcctccat |
| ***Mouse genotyping primers*** | |
| HLA-DR4-DQ8-F | tcccttgatgatgaagatgg |

| | |
|---|---|
| HLA-DR4-DQ8-R | cagaggtaactgtgctcacg |
| hCD4-F | ctttccagaaggcctccagca |
| hCD4-R | ctctcatcaccaccaggttcac |

[1]HKU16Δ*ssa* genomic DNA served as template and HKU16Δ*ssa* was used for genetic manipulation.

### References


1 Loh, J. M. & Proft, T. Toxin-antitoxin-stabilized reporter plasmids for biophotonic imaging of group A *Streptococcus*. *Appl Microbiol Biotechnol* **97**, 9737-9745 (2013)

2 Ly, D. *et al.* Plasmin(ogen) acquisition by group A *Streptococcus* protects against C3b-mediated neutrophil killing. *J Innate Immun* **6**, 240-250 (2014)

3 Sundberg, E. & Jardetzky, T. S. Structural basis for HLA-DQ binding by the streptococcal superantigen SSA. *Nat Struct Biol* **6**, 123-129 (1999)

4 Lee, K. H., Wucherpfennig, K. W. & Wiley, D. C. Structure of a human insulin peptide-HLA-DQ8 complex and susceptibility to type 1 diabetes. *Nat Immunol* **2**, 501-507 (2001)

5 Jardetzky, T. S. *et al.* Three-dimensional structure of a human class II histocompatibility molecule complexed with superantigen. *Nature* **368**, 711-718 (1994)

6 Wertman, K. F., Wyman, A. R. & Botstein, D. Host/vector interactions which affect the viability of recombinant phage lambda clones. *Gene* **49**, 253-262 (1986)

7 Tse, H. *et al.* Molecular characterization of the 2011 Hong Kong scarlet fever outbreak. *J Infect Dis* **206**, 341-351 (2012)

8 Timmer, A. M. *et al.* Streptolysin O promotes group A *Streptococcus* immune evasion by accelerated macrophage apoptosis. *J Biol Chem* **284**, 862-871 (2009)

9 Rivera-Hernandez, T. *et al.* Differing efficacies of lead group A streptococcal vaccine candidates and full-length M protein in cutaneous and invasive disease models. *mBio* **7**, pii: e00618-16 (2016)

10 Kasper, K. J. *et al.* Bacterial superantigens promote acute nasopharyngeal infection by *Streptococcus pyogenes* in a human MHC Class II-dependent manner. *PLoS Pathog* **10**, e1004155-e1004155 (2014)

11 Barnett, T. C., Daw, J. N., Walker, M. J. & Brouwer, S. Genetic manipulation of group A *Streptococcus* – gene deletion by allelic replacement. *In: T. Proft and J. Loh (Ed) Group A Streptococcus: Methods and Protocols. Springer, Heidelberg.*, in press (2020)
